# Could TDX-1 explain the changes observed in endocrine pancreas due to chronic cola drink consumption in rats? A global point of view

**DOI:** 10.1101/2020.11.20.391060

**Authors:** Gabriel Cao, Julián González, Juan P. Ortiz Fragola, Angélica Muller, Mariano Tumarkin, Marisa Moriondo, Francisco Azzato, Manuel Vazquez Blanco, José Milei

## Abstract

In previous studies, we reported evidence showing that chronic cola consumption in rats impairs pancreatic metabolism of insulin and glucagon and produces some alterations typically observed in the metabolic syndrome (i.e, hyperglycemia, and hypertriglyceridemia) with an increase in oxidative stress. Of note, no apoptosis nor proliferation of islet cells could be demonstrated. In the present study, 36 male Wistar rats were divided in three groups to freely drink regular cola, light cola, or water (controls). We assessed the impact of the three different beverages in glucose tolerance, lipid levels, creatinine levels and immunohistochemical changes addressed for the expression of insulin, glucagon, PDX-1 and NGN3 in islet cells, to evaluate the possible participation of PDX-1 in the changes observed in α and β cells after 6 months of treatment. On the other hand, we assessed by stereological methods, the mean volume of islets (Vi) and three important variables, the fractional β-cell area, the cross-sectional area of alpha (A α-cell) and beta cells (A β-cell), and the number of β and α cell per body weight.

Cola drinking caused impaired glucose tolerance as well as fasting hyperglycemia and increase of insulin immunolabeling. Immunohistochemical expression for PDX-1 was significantly high in regular cola consumption group compared to control. In this case, we observed cytoplasmatic and nuclear localization. Likewise, a mild but significant decrease of Vi was detected after 6 months of cola drink consumption compared with control group. Also, we observed a significant decrease of fractional β cell area compared with control rats. Accordingly, a reduced mean value of islet α and β cell number per body weight compared to control was detected. Interestingly, consumption of light cola increased the Vi compared to control. In line with this, a decreased cross-sectional area of β-cells was observed after chronic consumption of both, regular and light cola, compared to controls. On the other hand, NGN3 was negative in all three groups. Our results support for the first time, the idea that TDX-1 plays a key role in the dynamics of the pancreatic islets after chronic consumption of sweetened beverages. The loss of islets cells might be attributed to autophagy, favored by the local metabolic conditions.

## Introduction

Consumption of sugary beverages has been associated with an increase in all-cause mortality among older adults [1], primarily through an increase in cardiovascular disease associated mortality [2]. Currently, an estimated 1 of 6 deaths in the United States is attributed to coronary heart disease (CHD) [3]. In multiple experimental and long-term prospective studies, high consumption of dietary sugars, sugar-sweetened beverages (SSBs) in particular, has been associated with several CHD risk factors, including dyslipidemia [4], diabetes [5], and obesity [5,6].

Of note, **s**ugar-sweetened beverages are the single largest source of added sugar in the US diet^7^. Although consumption of SSBs has decreased in the past decade [8], data show a slight rebound in consumption in recent years among adults in most age groups. In developing countries, intake of SSBs is rising dramatically because of widespread urbanization and beverage marketing [9].

In previous studies, we reported evidence showing that chronic cola consumption in rats impairs pancreatic storage of insulin and glucagon and produces some alterations related to metabolic syndrome (hyperglycemia and hypertriglyceridemia) with an increase in oxidative stress [10–13]. Moreover, we also demonstrated that consumption of cola beverages, regardless sugar content, increased the rate of atherosclerosis progression in ApoE2/2 mice, favoring aortic plaque enlargement (inward remodeling) over media thinning [14,15].

Pancreatic duodenal homeobox 1 gene (Pdx1) encodes a transcription factor that critically regulates early pancreas formation and multiple aspects of mature beta cell function, including insulin secretion, mitochondrial metabolism, and cell survival. Reduced Pdx1 expression in the beta cell occurs in cellular models of glucose toxicity and accompanies the development of diabetes, correlating low Pdx1 levels with beta cell metabolic failure. Beside these aspects, little is known about the pathophysiologic process that justify the changes described in pancreatic islets by chronic consumption of SSBs in rats.

In this paper we focused the study on the effects in glucose and lipid homeostasis, pancreatic islets morphology and immunophenotype changes in alpha and beta cells observed after 6 months of chronic consumption of cola drink in male Wistar rats.

## Materials and methods

Animals were housed at the ININCA facilities (21±2°C, at 12-h light-dark cycles 7am-7pm) and were fed a commercial chow (16%-18% protein, 0.2 g % sodium (Cooperación, Buenos Aires, Argentina) *ad libitum*. Animal handling, maintenance and euthanasia procedures were performed according with international recommendations^16^. (Weatherall report, “The use of non-human primates in research.”). The committee of Ethics in Animal Research of the Instituto de Investigaciones Cardiológicas (ININCA) and the Institutional Animal Care and Use Committee (IACUC) of the Faculty of Medicine of the University of Buenos Aires (CICUAL, Institutional Committee for the Care and Use of Laboratory Animals) approved the study.

### Experimental protocol

Thirty-six male Wistar rats were randomly distributed in 3 groups (12 animals each group), which were respectively assigned to different treatments according to beverage (as the only liquid source, *ad libitum*): W (water), regular cola (C) (commercially available sucrose-sweetened carbonated drink, Coca-Cola, Argentina) and Light Cola (L) (commercially available low calorie aspartame-sweetened carbonated drink, Coca-Cola Light, Argentina) for six months (end of treatment). Rats were weighed weekly. Food and drink consumption were assessed twice a week. Biochemical assays were performed at baseline; biochemical assays and histopathological data from autopsies were obtained at the end of treatment (6 months).

According to company specifications, Coca Cola™ is a carbonated water solution containing (approximate %): 10.6 g carbohydrates, sodium 7 mg, caffeine 11.5 mg, caramel, phosphoric acid, citric acid, vanilla extract, natural flavorings (orange, lemon, nutmeg, cinnamon, coriander, etc), lime juice and fluid extract of coca (*Erythroxylum novogranatense*). As far as nutritional information is concerned the only difference between regular and light cola is the replacement of carbohydrates with non-nutritive sweeteners (aspartame + acesulfame K) in the latter [14]. Soft drinks had carbon dioxide content largely removed by vigorous stirring using a stirring plate and placing a magnetic bar in a container filled with the liquid prior to being offered to the animals at room temperature.

### Biochemical determinations

Plasma aliquots of blood collected from the tail vein after 4-hour fasting were used to measure the concentration of glucose and triglycerides by enzymatic colorimetric assays using commercially available kits (Sigma-Aldrich, USA) according to manufacturer’s instruction [17]. An oral glucose tolerance test^18^ was performed on each animal before and after treatment and plasmatic HDL and TG were also assessed. After treatment plasmatic urea, creatinine, ASAT and ALAT were measured.

### Immunohistochemistry

For immunohistochemical techniques, additional sections were cutting and mount in positively charged slides. Alpha and beta pancreatic cells were evaluated in dewaxed sections using specific primary antibodies (mouse monoclonal anti-Glucagon and anti-Insulin antibodies, Sigma-Aldrich Corp., St. Louis, Mo. USA). Rabbit polyclonal antibodies against PDX1, (pancreatic and duodenal homebox 1 also known as insulin promoter factor 1, ab47267, Abcam, Cambridge, UK), was used to evaluate the glucose dependent insulin transcription and its Type 1 diabetes association; and NGN3 (Neurogenin 3, ab176124, Abcam, Cambridge, UK), was used to assess α and β cells expression of endocrine pancreas and its link to neurogenesis.

Before staining, sections were deparaffinized and incubated in 3% hydrogen peroxide for 10 min to quench endogenous peroxidase. After washing 3 times in PBS, nonspecific antibody binding sites were blocked with 4% skimmed milk and 1% bovine albumin in PBS. Sections were incubated with the primary antibodies diluted in blocking solution at 4°C overnight. Negative controls were incubated with 4% skimmed milk and 1% bovine albumin in PBS. Sections were then washed 3 times in PBS and subsequently incubated with a biotinylated secondary anti-mouse or anti-rabbit antibody diluted (provided by Ch. Lillig, Germany) for 60 min at room temperature. Immunohistochemical staining was obtained performed using a biotinylated-streptovidin-peroxidase complex (Dako Universal LSAB™+ Kit/HRP-K0690, Dako, Glostrup, Denmark) with DAB (3,3-Diaminobenzidin)-chromogen (Dako-K3468, Dako, Glostrup, Denmark) as detection system according to manufacturer recommendations.

### Morphologic Analysis

At 6 months, necropsy was practiced. The whole pancreas was weighed and fixed in buffered 10% formaldehyde solution for 24h at room temperature. Later, the tissues were dehydrated in alcohols, cleared in xylene and embedded in paraffin. For light microscopy, a Nikon Eclipse 50i microscope (Nikon Corporation, Tokyo, Japan), equipped with a digital camera (Nikon Coolpix S4) and the Image-Pro Plus image processing software version 6.0 (Media Cybernetics, Silver Spring, Maryland, USA) were used. For stereological analysis, 3 μm width sections were cut from tissue blocks and stained with hematoxylin-eosin. Thus, digitalized images of 12 to 16 pancreatic islets per animal, were obtained by systematic uniform random sampling of the tissue (total islets evaluated per group: 144 – 192). At this time, an orthogonal grid with 50 test points, representing a total area of 6.7 10^4^μm^2^ (objective lens magnification: 40 X) was employed. Points were projected onto the fields of view and the number of points hitting the pancreatic islets was counted. In this way, the Cavalieri method with point-counting [19] was employed to estimate the mean volume of pancreatic islets:

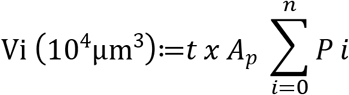

*t*: mean section cut thickness; *A_p_*: area associated with a point.

*P_i_:* points counted on grid.

The evaluation of immunolabeled slides was carried out using a methodology that include the transformation of the digitized images to the CMYK color scheme, using Adobe Photoshop v 21.0.2 software [20]. Subsequently, the images were separated into color channels, selecting the yellow channel image for analysis, previously converted to gray scale image. Finally, with appropriately calibrated image analysis software, the integrated optical density (IOD) of the immunolabelled insular areas was estimated. The IOD results from the product of the optical density (OD) and the immunolabeled tissular surface.

For glucagon:

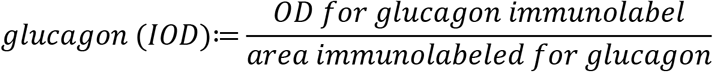

For insulin:

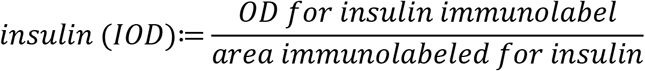

For TDX-1:

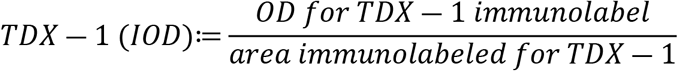

For NGN3:

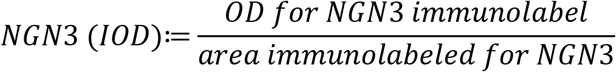

The fractional beta cell area (Faβ) is an important and predictive parameter, given that their reduction is linearly related to insulin resistance [21]. Also, is directly influenced by the beta cell mass changes. Usually is estimated by the quotient between the islet immunolabelled area for insulin and the pancreatic tissue surface, including the exocrine pancreas. From our point of view, it is more appropriate to use the area of the islet as denominator, excluding the pancreatic glandular component:

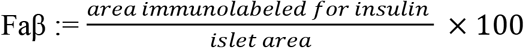

In this way, Faβ was estimated by point counting, representing the mean area of pancreatic islet immunomarker for insulin and expressed in percent (%).

On the other hand, the number of alpha (nα) and beta (nβ) cells in the studying islets were estimated from the insular area immunolabeled for glucagon and insulin respectively, and the cross-sectional area of individual nucleated cell per islet. Cross-sectional area was carried out with the image processing software at 400X, considering spherical the cellular shape. We take the maximal diameter of each insular cell and calculate their area by applicate the surface area of sphere formula. A sphere with radius r has a volume of:

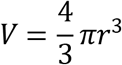

then, their corresponding surface is:

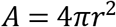

Given the relationship between body weight (BW) and nβ [22], we present this variable as the quotient between nα, nβ and BW respectively (nα g^−1^ and nβ g^−1^):

For alpha cells:

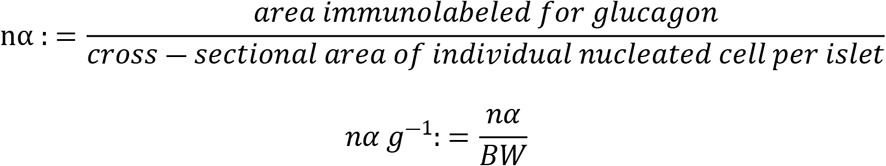

Cross-sectional area of individual nucleated islet cell represents the area of alpha and beta cells employed to analyze the structural changes observed along the experiment.

For beta cells:

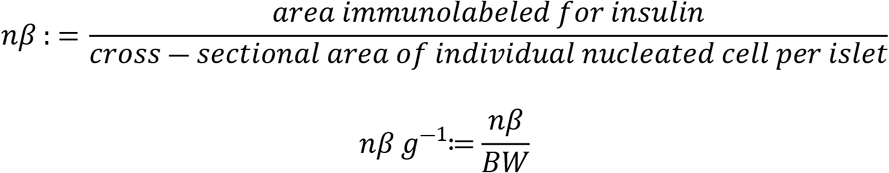

### Statistical analysis

Data were analyzed by two-way ANOVAs followed by post-hoc tests (Bonferroni multiple t-test) or by Kruscal-Wallis followed by Dunn test (depending on distribution) in order to evaluate between-groups’ differences. Statistical significance was set at p<0.05 and SPSS version 15.0 software was used to analyze data.

## Results

### Nutritional considerations

Regular cola drinking for 6 months caused an increase in liquid intake and a decrease in food intake (p < 0.05). Conversely, Light cola drinking resulted in an increase in food and liquid intake (p< 0.05). Body weight increased with no significant differences between groups (Table 1).

**Table 1.** Nutritional aspects and plasmatic parameters after treatment. Treatment: water (W), regular (C), or light cola drinking (L). * p<0.05 vs W.

| Variable | W | C | L |
| --- | --- | --- | --- |
| Solid intake (g/kg/day) | 43±4 | 31±3.7* | 46±3.8* |
| Liquid intake (ml/kg/day) | 74.1±13.1 | 138±33* | 92.8±26.4* |
| Body weight (at 6 months) | 119.7±3.1 | 118.8±4.4 | 116.8±9.7 |
| Creatinine | 0.53(0.51-0.54) | 0.56(0.55-0.61)* | 0.6(0.55-0.63)* |
| ASAT | 69(54-87) | 70(52-109) | 81(60-108) |
| ALAT | 59(47-62) | 63(42-73) | 55(46-87) |

### Biochemistry

No significant differences were found at baseline (before randomization) between groups. Compared to control, regular cola drinking for 6 months produced alterations on lipid and glucose homeostasis and on renal function (p<0.05). Glycemia was significantly higher in cola drinking rats both fasting, at 30 and at 120 minutes (Table 2). On the other hand, HDL was significantly diminished. Triacylglycerols were markedly higher in the cola group, but it did not reach statistical significance, perhaps due to great data dispersion. Creatinine was also elevated in cola drinking rats (p<0.05) (Table 1). Light cola drinking resulted in no significant changes in glycemia and lipid profile, but also increased creatinine (p<0.05) (Table 2).

**Table 2.** Metabolic parameters before and after treatment. Delta: 6 months-baseline difference. Oral glucose tolerance test: Glu 1: basal glucose, Glu 2: glucose after 30 minutes. Glu 3: final glucose. HDL: high density lipoprotein. TG: triglycerides. Treatment: water (W), regular (C), or light cola drinking (L). * p<0.05 vs W.

| Variable | Baseline |  |  | 6 months |  |  | Delta |  |  |
| --- | --- | --- | --- | --- | --- | --- | --- | --- | --- |
|  | W | C | L | W | C | L | W | C | L |
| Glu 1 | 137(101-158) | 122.5(116-143) | 127(121-133) | 125(120-129) | 131(124-138)* | 125(116-129) | -12(-35-18) | 6(-8-14) | -7(-25-9) |
| Glu 2 | 183(179-211) | 181(169-203) | 178(170-197) | 145(139-154) | 157(152-160)* | 136(128-143)* | -45(-63-25) | -31(-43--12.5)* | -45(-75--38) |
| Glu 3 | 131(119-156) | 120(109-139) | 126(119-142) | 135(129-142) | 148(137-153)* | 131(121-133) | 0(-23-16) | 22(16-34)* | 4(-5-10) |
| HDL | 21(17-27) | 22(21-24) | 21(18-21) | 33(30-36) | 25(22-31)* | 28(22-49) | 11(9-18) | 3,5(-0.5-9)* | 8(4-28) |
| TG | 70(56-94) | 64(55-78) | 60(40-74) | 138(132-172) | 258(106-479) | 111(97-403) | 73(48-83) | 200(45-397) | 67(34-329) |

### Morphology and Immunohistochemistry

After 6 months of treatment, C group showed significant increase of insulin IOD compared to W and L groups (p<0.0001, respectively). Conversely, not significant differences were seen between L and W groups. On the other hand, cytoplasmatic expression for glucagon in C group was highest than W and L (p<0,0001, respectively). Interesting, L group showed lowest glucagon IOD compared to W and C (p<0.0001, respectively). Consumption of regular cola for 6 months significantly reduced the nuclear and/or cytoplasmic expression of PDX-1 in male Wistar rats, compared to W and L groups (both p<0.0001). Similar values of PDX-1 IOD were seen in W and L (see Table 3 and Figure 1).

**Table 3.**
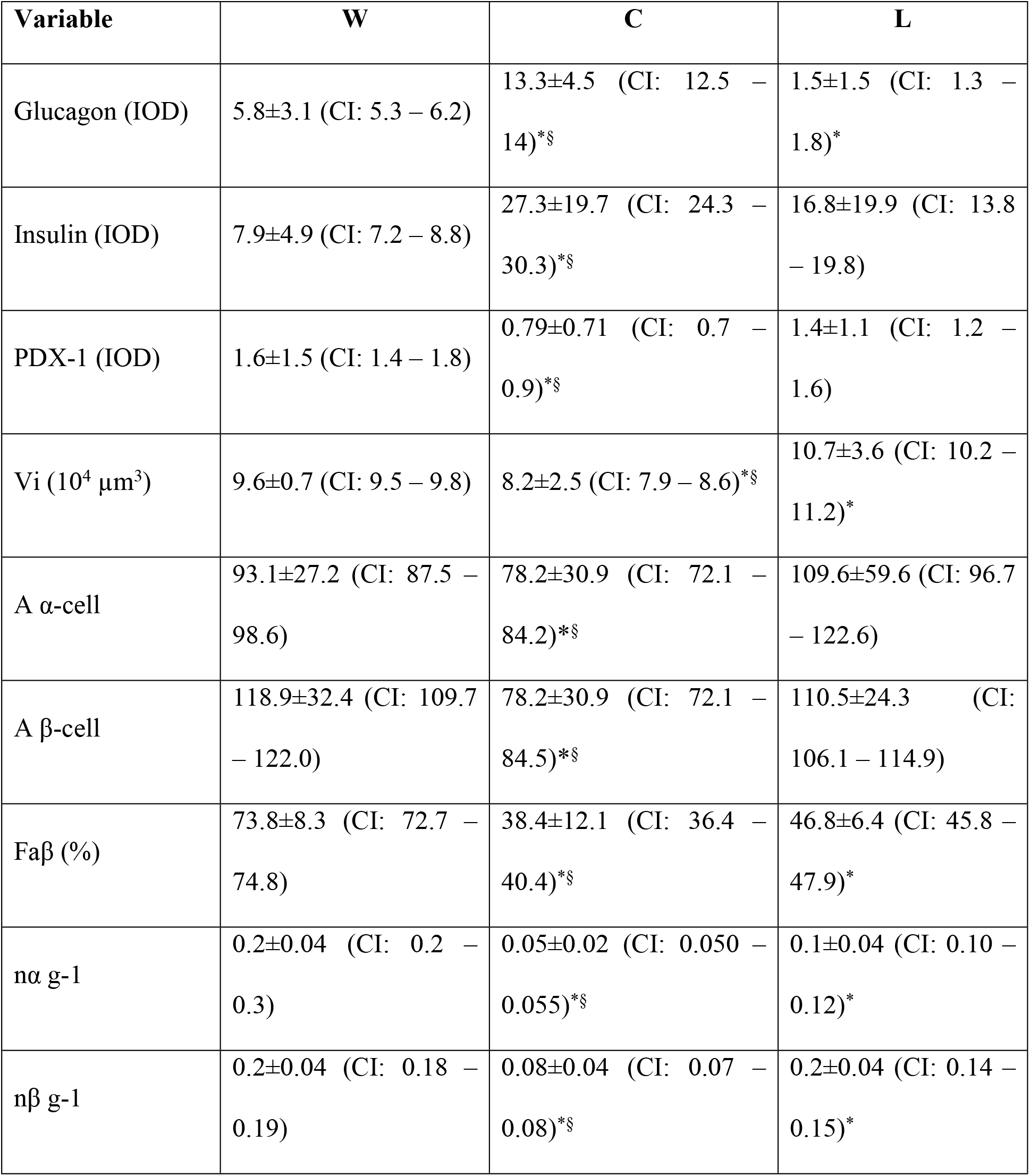
Immunohistochemistry comparisons for insulin, glucagon and PDX-1 after 6 months of treatment. Also, we show stereological variables, such as: Vi (10^4^ μm^3^) (mean pancreatic islet volume), A α-cell and A β-cell (cross-sectional area of α and β cells), Faβ (%) (fractional β cell area), nα g-1 and nβ g-1 (mean value of islet α and β cell number per BW). Treatment: water (W), regular (C), or light cola drinking (L). * p<0.05 vs W; § p< 0.05 vs L.

| Variable | W | C | L |
| --- | --- | --- | --- |
| Glucagon (IOD) | 5.8±3.1 (CI: 5.3 – 6.2) | 13.3±4.5 (CI: 12.5 – 14)*§ | 1.5±1.5 (CI: 1.3 – 1.8)* |
| Insulin (IOD) | 7.9±4.9 (CI: 7.2 – 8.8) | 27.3±19.7 (CI: 24.3 – 30.3)*§ | 16.8±19.9 (CI: 13.8 – 19.8) |
| PDX-1 (IOD) | 1.6±1.5 (CI: 1.4 – 1.8) | 0.79±0.71 (CI: 0.7 – 0.9)*§ | 1.4±1.1 (CI: 1.2 – 1.6) |
| Vi (10<sup>4</sup> µm<sup>3</sup>) | 9.6±0.7 (CI: 9.5 – 9.8) | 8.2±2.5 (CI: 7.9 – 8.6)*§ | 10.7±3.6 (CI: 10.2 – 11.2)* |
| A α-cell | 93.1±27.2 (CI: 87.5 – 98.6) | 78.2±30.9 (CI: 72.1 – 84.2)*§ | 109.6±59.6 (CI: 96.7 – 122.6) |
| A β-cell | 118.9±32.4 (CI: 109.7 – 122.0) | 78.2±30.9 (CI: 72.1 – 84.5)*§ | 110.5±24.3 (CI: 106.1 – 114.9) |
| Faβ (%) | 73.8±8.3 (CI: 72.7 – 74.8) | 38.4±12.1 (CI: 36.4 – 40.4)*§ | 46.8±6.4 (CI: 45.8 – 47.9)* |
| nα g-1 | 0.2±0.04 (CI: 0.2 – 0.3) | 0.05±0.02 (CI: 0.050 – 0.055)*§ | 0.1±0.04 (CI: 0.10 – 0.12)* |
| nβ g-1 | 0.2±0.04 (CI: 0.18 – 0.19) | 0.08±0.04 (CI: 0.07 – 0.08)*§ | 0.2±0.04 (CI: 0.14 – 0.15)* |

**Figure 1.**
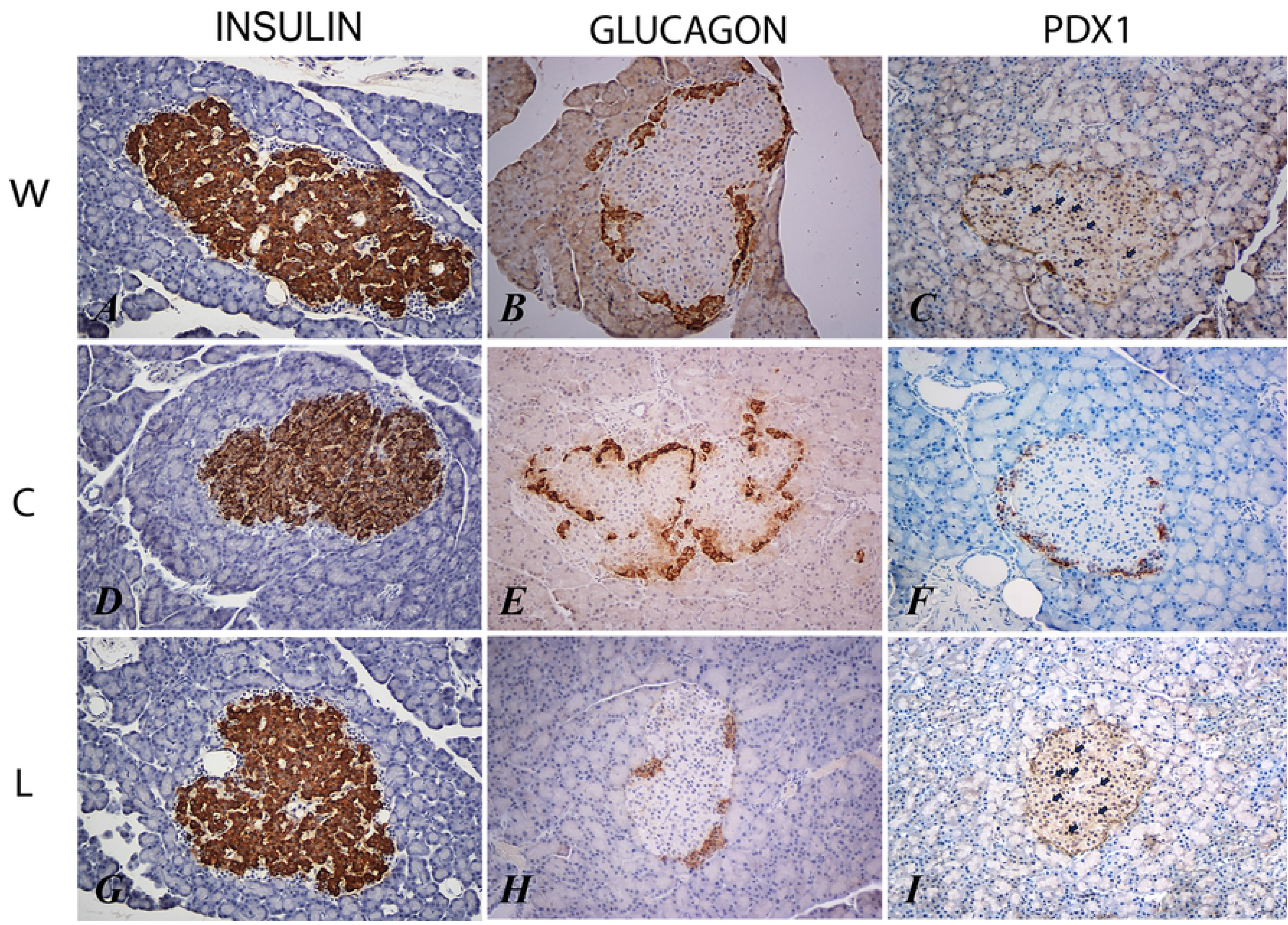
Immunostaining for insulin (A, D, G), glucagon (B, E, H) and PDX-1 (C, F, I) in water drinking (W), cola drinking (C), and light cola drinking (L) experimental groups (see text for details). In C and I, black arrows indicate nuclear immunostaining for TDX-1.

Significant reduction of Vi were seen in C group, compared to W (p<0.0006) and L (p<0.0001). Conversely, L group showed a significant increase of Vi compared to W (p<0.0006). Reductions of Faβ were seen in C and L groups compared with W (p<0.0001, respectively). Likewise, a lowest Faβ value was seen in C than L group (p<0.0001). Similar profile was observed in C and L groups for nα g^−1^ and nβ g^−1^ values, compared to W group (p<0.0001, respectively). Also, C demonstrated lowest mean values than L group (p<0.0001, in both cases). Finally, NGN3 was negative in all three groups (see Table 3; Figures 2 and 3).

**Figure 2.**
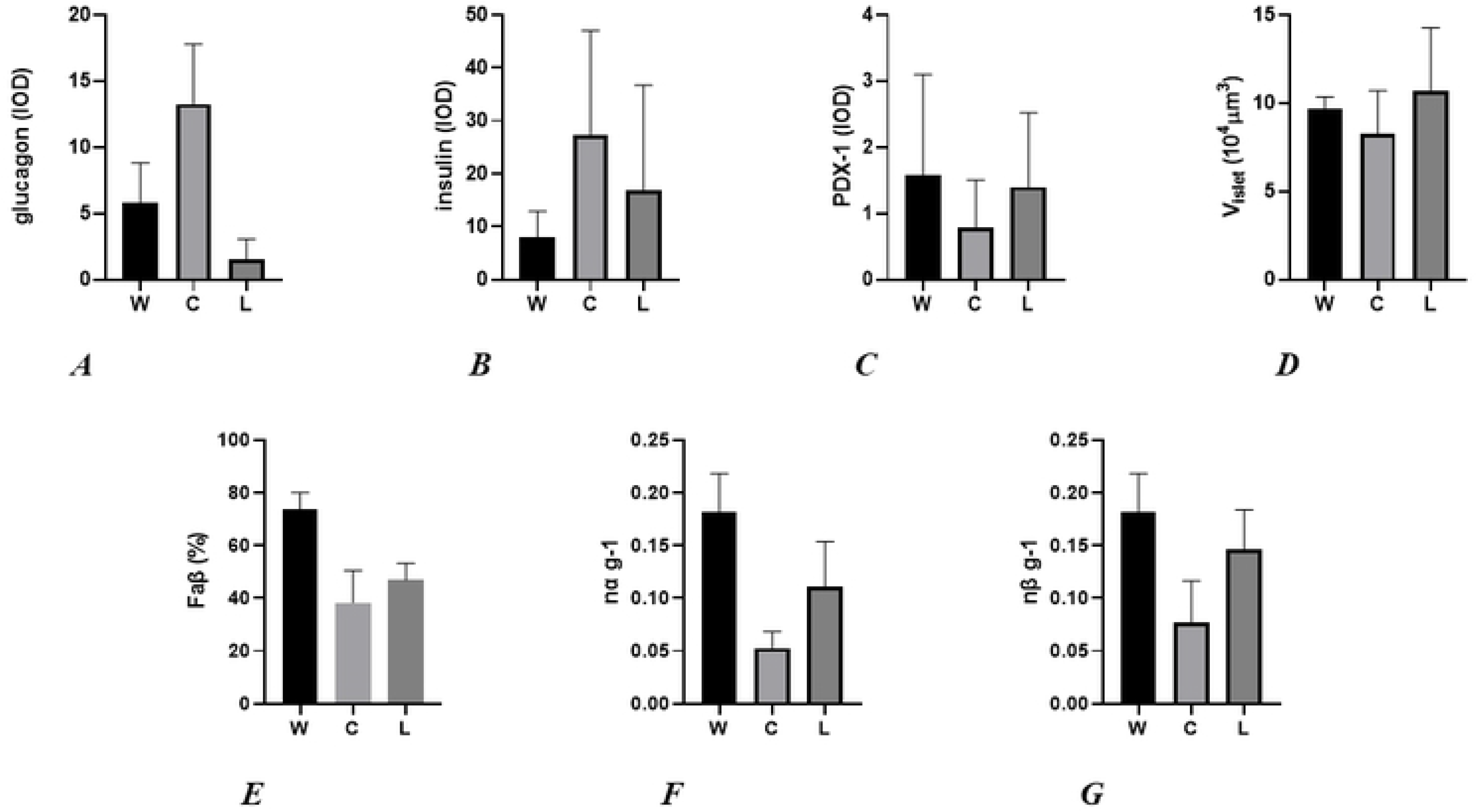
Graphics’ A, B, C: IOD for glucagon, insulin, and PDX-1 respectively; D – G: mean pancreatic islet volume, fractional β cell area and mean value of islet α and β cell number per BW in water drinking (W), cola drinking (C), and light cola drinking (L) experimental groups.

**Figure 3.**
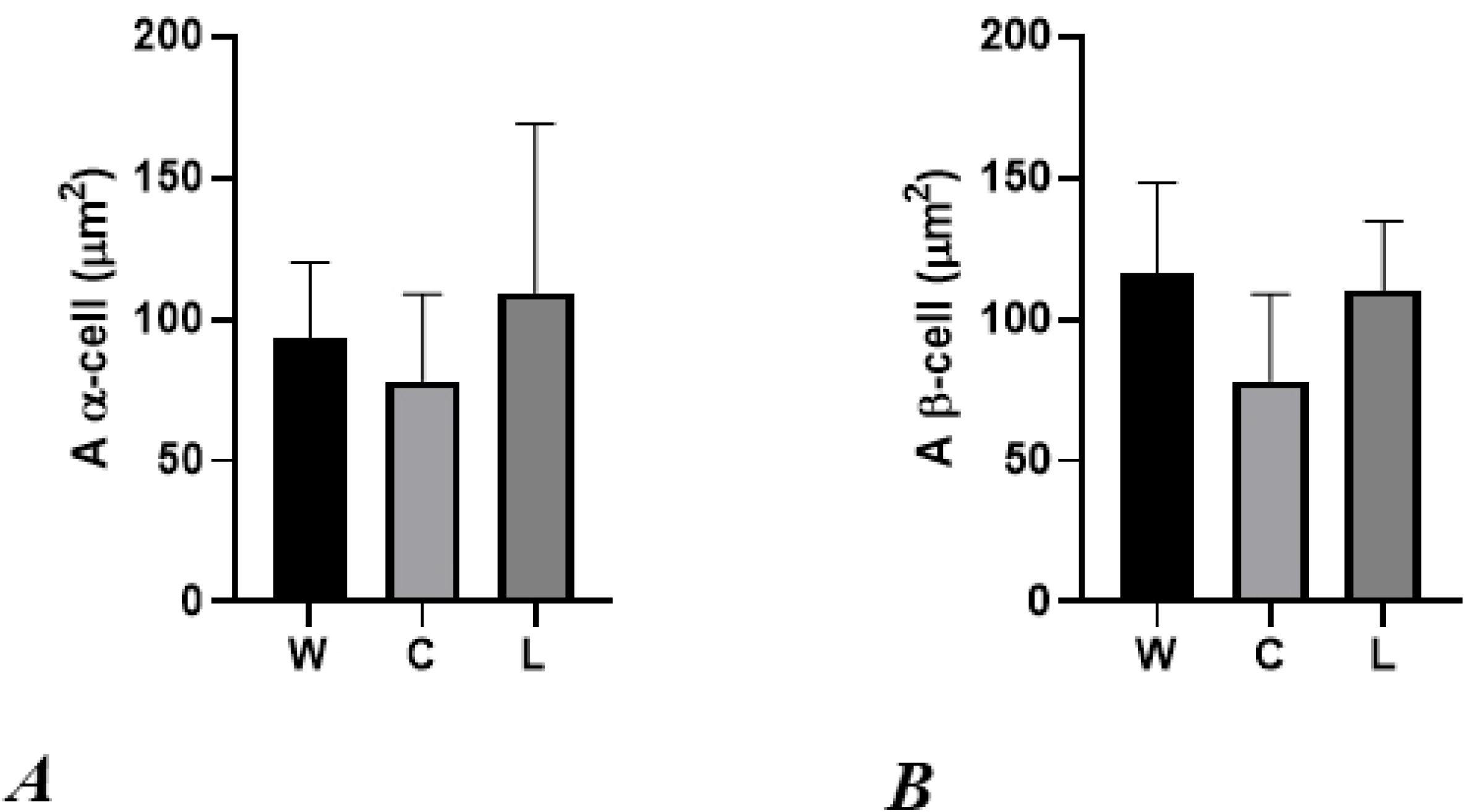
Graphics’ A and B represent the cross-sectional area of α and β cells respectively in water drinking (W), cola drinking (C), and light cola drinking (L) experimental groups.

## Discussion

In this paper, we show that cola drinking for 6 months produced alterations on lipid and glucose homeostasis, in addition to changes in the structure and immunolabeling profile of pancreatic islets. Regular cola drinking caused an increase in liquid intake (empty calories) and a decrease in food intake, while light cola drinking resulted in an increase in food and liquid intake; none of these effects were associated with changes in body weight. These results are similar to our previous studies [12,13,23], and studies by others [24–26]. The increase in fluid intake is partially explained by sweet taste and sugar content rather than increased thirst. This may be related to an “hedonic” effect caused by sugar ingestion. Excessive sugar consumption is related to decreased striatal concentrations of dopamine [27]. On the opposite, acute effects of sugar consumption increase dopamine in the nucleus *accumbens*, an area closely related to reward feelings [28]. Some experiments with opioid-receptor knockout mice suggest that opioid system may also be involved in reward sensation related to sugar intake [29,30]. In this connection, behavioral studies in mice showed a preference for caloric liquids independently of sweet taste [31].

The effect on solid food consumption of regular and light cola drinks may be related also to reward. The effect of sucrose on appetite is complex. In a mice study, animals given a premeal load of sugar consumed more chow. In the same study, sugar seemed to decrease the content of neuropeptide Y and agouty-related protein in the arcuate nucleus, both with orexigenic effect, immediately after sugar ingestion, but then showed to cause a sort of rebound effect with an increase of both neurotransmitters just before chow ingestion [32]. Besides, the effect of glucose and fructose (both components of sucrose) seem to be opposite. While glucose leads to an increase in the hypothalamus content of cholecystokinin (satiety-inducing) [33] and decreased ghrelin (orexigenic) [31], fructose makes the contrary [30,32]. On the other side, both chronic consumption of glucose and fructose decrease polypeptide YY serum levels and hypothalamic proopiomelanocortin mRNA expression, both with satiety-inducing effects [34]. Taking all these data into account, sugar should prompt an increase in food intake, but the opposite is observed very consistently in the many studies cited [10,12,13,23–25,32,33,35].

The role of caffeine has also been matter of debate in some cola drinking studies. Caffeine has shown to enhance thermogenesis and energy expenditure in rats [36], but doses used in this paper are far higher than doses we calculated to be provided by cola and light cola drinking [12]. Consistently with other studies, we found that regular cola impaired glucose tolerance and decreased HDL cholesterol [10–12,24]. This hyperglycemic effect is explained by several mechanisms, including impairment of the expression of adipocytokines, altered hepatic glucose management with increased glycogenolysis, gluconeogenesis and pancreatic dysfunction [2,10].

Regular cola drinking also induced an increase in triacylglycerols levels. This can be ascribed to the excess of calories delivered by sugar content of cola beverages and specially to fructose content. Glucose enter glycolytic pathway and is metabolized to pyruvate, which in turn is converted to acetyl-CoA, which then enters de tricarboxylic acid cycle. When glucose is abundant in hepatocytes, tricarboxylic acid cycle saturates, and excess of acetyl-CoA is derived to fatty acid synthesis and then to triacylglycerols synthesis. Fructose enters the hepatocyte and is phosphorylated in carbon 1, bypassing the limiting enzyme of glycolysis, phosphofructokinase-1^40^. That explains the relation between fructose consumption and fatty liver disease [40].

After 6 month of regular cola consumption, pancreatic islets showed a significant immunolabeling increase for insulin in both C and L groups. In light cola group, this phenomenon might be explained by two mechanisms: 1-acesulfame K and other non-nutritive sweeteners can activate incretin release and stimulate pancreatic insulin production [37,38], and 2-phenylalanine, partially metabolized to aspartame, can synergistically stimulate insulin secretion [38–40]. This mechanism could be justifying the glycemia decrease observed in the glucose tolerance test in light cola drinkers (improved glucose tolerance).

Long time consumption of regular cola as unique source of liquid induced a significant increase of glucagon immunolabeling compared with W group. On the other side, in the same period, consumption of cola light suppressed the immunolabeling of glucagon to values lowest than W and C groups. In rats, mechanisms associated to stimulation or inhibition of glucagon release includes hyperglycemia and hypertriglyceridemia, respectively. Some reports show that blood glucose concentration from 7 to 8 mmol/L, maximally inhibit glucagon secretion [41,42]. Interestingly, concentrations above 20 mmol/L stimulate glucagon secretion in mouse [43]. This phenomenon was associated with variation of cytosolic levels of Ca^2+^. Also, hypertriglyceridemia and increased blood fatty acid levels reduce glucagon release in rats, dogs and guinea pigs [44]. Likewise, chronic consumption of acesulfame K inhibit glucagon synthesis by stimulation of glucagon-like peptide-1 (GLP-1), an incretin produced in the gut [45,46]. Also, long time consumption of acesulfame K and aspartame increase blood levels of triglycerides, ASAT and ALAT transaminases, creatinine and urea, reducing blood levels of HDL cholesterol [46]. Finally, in C group, high production of insulin hypothetically could inhibit release of glucagon through the production of somatostatin from δ cells. Nevertheless, low levels of glucagon persist, suggesting that hyperglycemia possess greater metabolic influence on α cells (Fig. 1).

Chronic consumption of regular cola, reduced the mean islet Vi, associated to increased immunolabeling for insulin, compared to W group. This stereological change could be explained by reduction of area and number of β-cells, involving an increased synthesis and release for insulin each individual cell. This finding suggests develop of metabolic stress. In previous papers [10,13] we reported an increased expression of insular thioredoxin-1 and peroxiredoxin-1 in cola drinking rats, suggesting a key participation of oxidative stress in the pathogenic changes observed in pancreatic islets. This scenario inhibits PDX-1 activity and critically suppress the cellular capability for hypertrophy, hyperplasia and protection of β-cells [47–49]. In previous papers [10,13] we tested the participation of apoptosis and proliferation related to the structural changes observed in pancreatic islets, by immunostaining for caspase 3 and PCNA [10]. Low expression was seen, associated to scarce insular PDX-1 activity, especially in C group. Also, oxidative stress induces the nucleo-cytoplasmic translocation of PDX-1, justifying the cytoplasmic localization observed by immunohistochemical studies [50]. The changes described above lead to significant reduction of Faβ, indicative of peripheral insulin resistance. Persistence of this anomaly, maintains insulin “hyperproduction” worsening the paracrine autoregulation of the pancreatic islet, conducing finally to “dysregulation” of this structure, condition frequently seen in type 2 diabetic patients [51].

In this point, we note the logical relation between oxidative stress, reductions in Vi, cross-sectional area and number of β-cells and the lowest proliferative and apoptotic activity. Accumulation of reactive oxygen species (ROS) and reactive nitrogen species (RNS) being among the main intracellular signal transducers sustaining autophagy. We postulate that long time consumption of regular cola in rats increase oxidative stress in pancreatic islets, inducing cellular loss via autophagy, independent of caspase cascade. In turn, size reduction of β-cells in C group compared with W group, could be explained by the metabolic demands, the low levels of ATP and the cellular need to obtain molecules and energy, which is obtained from own cellular components. Consumption of light cola for 6 months provoked a modest but significantly increased mean Vi value compared to W and C groups. Also, both nα g^−1^ and nβ g^−1^ demonstrated reduced values compared to W group, but to minor severity than C group. Interestingly, a non-significantly increased value of cross-sectional area for α cells was seen, compared to W group (Fig. 3). Although, taking into account the values dispersion, we clearly noted that mean nominal value of A α cell was increased, suggesting a possible influence on the mean increased value of Vi, given that not significantly changes were observed in cross-sectional areas of beta cell after chronic consumption of cola light compared to W group (Fig. 3). In a previous paper, our group could not demonstrate proliferative activity in pancreatic islets [10], thus this phenomenon may be attributed to mild α cell hypertrophy, associated to significantly reduction of glucagon IOD and possibly mediated by PDX-1 activity. In L group, variables such as Vi, nα g-1, nβ g-1, A α cell and A β cell (Fig. 2 and 3) showed similar profile than C group, but with minor severity. These finding could be related to the minor caloric delivery by light cola, contrarily to regular cola. In this sense, long term consumption of light cola potentially could produce also similar metabolic disturbances as observed in type 2 diabetes.

Consumption of light cola for 6 months provoked a modest but significantly increased of mean Vi value compared to W and C groups. Also, both nα g^−1^ and nβ g^−1^ demonstrated reduced values compared to W group, but to minor severity than C group. Interestingly, a non-significant increased value of cross-sectional area for α cells was observed, compared to W group (Fig. 3). Although, taking the values dispersion into account, we clearly note that mean nominal value of A α cell was increased, suggesting a possible influence on the mean increased value of Vi, given that not significant changes were observed in cross-sectional area of beta cell after chronic consumption of cola light compared to W group (Fig. 3). In a previous paper, our group could not demonstrate proliferative activity in pancreatic islets [10], thus this phenomenon could be attributed to mild α cell hypertrophy, associated to significantly reduction of glucagon IOD and possibly mediated by PDX-1 activity (Fig. 1). In L group, variables such as Vi, nα g-1, nβ g-1, A α cell and A β cell (Fig. 2 and 3) showed similar profile than C group, but with minor severity. These finding may be related to the minor caloric delivery by light cola, contrarily than regular cola. In this sense, long term consumption of light cola potentially may also produce similar metabolic disturbances such as observed in type 2 diabetes.

The transcriptional factor neurogenin 3 (NGN3) is involved in differentiation, transdifferentiation and regeneration of pancreatic islet cells [53]. We could not demonstrate NGN3 activation in pancreatic islets after chronic consumption of regular cola. Then, and in opposition to our previous hypothesis [10], transdifferentiation seems not to be the principal phenomenon that explains the dynamics of pancreatic islet cells, observed in this model.

## Conclusions

Consumption of regular cola after 6 months as unique source of liquid, induces local and systemic oxidative stress in rats. At pancreatic islet level, accumulations of ROS and RNS damage β cells, through PDX-1 activity suppression, promoting autophagic increase and consequent cellular loss. In this point, β cell promoting an increased compensatory synthesis, storage and release of insulin. This scenario favors metabolic stress and lack of ATP, accelerates cellular loss and islet dysfunction. These events conduct to Faβ decrease, an important morphologic variable associated to insulin resistance. This pathophysiologic mechanism explains pancreatic islet changes observed after 6 month of regular cola consumption (Fig. 2 and 3). On the other side, long term light cola consumption may develop a similar mechanism than regular cola, but for a longer time due to the lower caloric intake it provides.

Our results support for the first time, the idea that TDX-1 plays a key role in the dynamics of the pancreatic islets after chronic consumption of sweetened beverages. The loss of islets cells might be attributed to autophagy, favored by the local metabolic conditions (Fig. 2 and 3).

## Acknowledgments

We especially acknew to Nora Paglia, VMR, for the professional support and dedicated supervision of this experiment.

## Founding

This study was supported by UBACYT 20020150100027BA from University of Buenos Aires, Argentina.

